# Virtual Reality system for freely-moving rodents

**DOI:** 10.1101/161232

**Authors:** Nicholas A. Del Grosso, Justin J. Graboski, Weiwei Chen, Eduardo Blanco-Hernández, Anton Sirota

## Abstract

Spatial navigation, active sensing, and most cognitive functions rely on a tight link between motor output and sensory input. Virtual reality (VR) systems simulate the sensorimotor loop, allowing flexible manipulation of enriched sensory input. Conventional rodent VR systems provide 3D visual cues linked to restrained locomotion on a treadmill, leading to a mismatch between visual and most other sensory inputs, sensory-motor conflicts, as well as restricted naturalistic behavior. To rectify these limitations, we developed a VR system (ratCAVE) that provides realistic and low-latency visual feedback directly to head movements of completely unrestrained rodents. Immersed in this VR system, rats displayed naturalistic behavior by spontaneously interacting with and hugging virtual walls, exploring virtual objects, and avoiding virtual cliffs. We further illustrate the effect of ratCAVE-VR manipulation on hippocampal place fields. The newly-developed methodology enables a wide range of experiments involving flexible manipulation of visual feedback in freely-moving behaving animals.

## INTRODUCTION

Movement is a fundamental element in the action-perception loop that is critical for most cognitive functions, such as decision-making, memory and spatial navigation. Internally-driven locomotor, head and sensor movements, an exploratory repertoire of a naturally-behaving animal, allow it to actively sample sensory information from the outside world for its optimal detection and encoding, as well as guidance of the behavior^1-5^. Recognition that the closed-loop link between internal dynamics, motor output and sensory processing gives rise to predictive coding, attention and flexible motor control^6-8^ is encouraging the use of a new experimental paradigm in sensory and cognitive neuroscience: closed-loop sensory stimulation. Traditional open-loop experimental paradigms involving head-fixation of the animal, useful for performing sensitive measurements of functional brain activity, are being replaced by experimental setups that partially close the loop between action and sensation while still retaining precise control of sensory inputs^9-13^.

Virtual reality (VR) systems close the loop between locomotion and vision. Many rodent laboratories use head-or body-restrained VR (rVR) setups to simulate locomotion through a 3D virtual environment (VE) via running on a treadmill^10,11^. Spatial coding research has especially benefited from such systems; VR researchers have taken advantage of the flexibility of a VE by implementing arbitrarily-large environmental exploration paradigms utilizing dynamic environments^14-16^ and manipulating visuomotor gain^17^. Additionally, many researchers take advantage of the rodent's fixed head by performing optical and intracellular recordings during locomotion through virtual space, a normally-challenging task in freely-moving animals^10,18,19^.

However, locomotion on a treadmill alone may not be enough for performing closed-loop research; behavioral and physiological differences between rVR and real-world navigation illustrate the detrimental effect of sensorimotor loop disruption and the importance of increasing motor affordances. While head-fixed rodents in rVR experiments are limited to navigating linear tracks^10,17-20^, likely due to an impoverished sensory-motor loop (Schmidt-Hieber, personal communication), rodents can navigate a two-dimensional VE if only their bodies are restrained and their heads left free to move^14,16,21-23^. If rats are further allowed to rotate while running on a spherical treadmill in rVR experiments, 2D hippocampal place cell representation of the VE is comparable to that in real-world navigation^23^; however, this effect is lost if the rodent's body rotation range is limited^21,24^.

Despite the great utility of rVR for studies of spatial navigation, animal restraint still poses unresolved challenges. First, restrained animals exhibit constrained or limited behavioral patterns within 2D space, which affects the way they actively sample the 3D environment. Second, locomotion-driven visual input is in conflict with locomotion-independent, head-bound idiothetic, olfactory, tactile and auditory inputs. Third, proprioceptive and vestibular inputs in rVR setups are diminished and unnatural, making them potential causes of the observed reduction in frequency-and speed-correlates of theta oscillatory dynamics, compared to rodents allowed to freely navigate the real world^23,24^. Lastly, animals require long and complex training and habituation to rVR setups^23,25^.

These challenges are resolved if visual feedback in VR is based on head motion in 3D space in freely-moving subjects, giving rise to a coherent visual, idiothetic and external multisensory input, an unperturbed action-perception loop, and a full repertoire of rodent behavior, while still preserving the precise control of visual stimuli in VR setups^26^. One such freely-moving VR (fmVR) system was introduced for human subjects as the Cave Automatic Virtual Environment (CAVE)^27^. A CAVE allows observers to freely move in space and view a 3D VE on the projection surfaces surrounding them. To date, CAVE-like VR systems for flies^28,29^ and fish^30^ couple animal 3D motion to 2D contrast patterns on the projected onto cylindrical surfaces, though a system for arthopods with more realistic visual feedback was reported^31^. Implementation of the CAVE system in rodents, a model mammalian system where complex interrogation and manipulation of the nervous system can be combined with cognitive behavior, would open new dimensions in experimental neuroscience. Designing an immersive fmVR in quickly-moving animals is challenging, however, as it would require very-low-latency visual feedback to avoid introducing new conflicts in the sensorimotor loop^32^ and computationally-intensive graphical operations to produce a visually-rich VE. An urgent need for and benefits of the development of a next-generation immersive fmVR were called for in a recent review^11^.

To provide an immersive virtual environment for untrained freely-moving rodents and allow them to explore and interact with the virtual environment in a natural manner, we developed a new CAVE fmVR system (ratCAVE) that produces minimal inter-sensory conflict during self-motion using fast head-tracking and high display frame rates, as well as enriched visual 3D cues of the virtual scene. We demonstrate the naturalistic interaction of rats with VEs in our fmVR system in several behavioral tasks. We further show a use case of fmVR not possible with rVR systems: to study the multisensory nature of hippocampal spatial representation. This highly-immersive fmVR system can be a powerful tool for a broad range of neuroscience disciplines.

## RESULTS

### ratCAVE: VR system for freely moving rodents

We implemented a CAVE system where a VE projection on the surface of the arena was closed-loop coupled with the real-time tracking of the head of the animal. In this setup, animals could move freely in a rectangular arena similar to that used for conventional open-field experiments, but the white-painted arena served as a projection surface. We used an array of 12 high-speed cameras (240-360 fps, NaturalPoint Inc.) to track the 3D position of the rodent’s head via a rigid array of retro-reflective spheres attached to a head-mounted 3D-printed skeleton (Fig. 1c,d). This tracking system enabled us to update the rodent’s head position with very high spatial (<0.1 mm) and temporal (<2.7 msec) resolution. The VE, created using open-source 3D modeling software (Blender 3D), was rendered each frame in a full 360-degree arc about the rodent’s head and mapped onto a 3D computer model of the arena using custom Python and OpenGL packages (Supplementary Fig.3, Online Methods), warped in real-time to generate a fully-interactive, geometrically-accurate 3D scene (Fig. 1b). The core cube-mapping algorithm used to perform the mapping of the VE onto the projection surface was identical to those described in rodent rVR setups (Supplementary Fig. 2a-c)^23^, but VE projection onto the surface of the arena is continuously updated according to the changing 3D position of the rodent’s head (Fig 1b), resulting in perception of a 3D VE that is stable in the real-world frame of reference that the animal is freely moving about (Fig 1c,d). The resulting image was front-projected onto the floor and slanted walls of the arena from a ceiling-mounted high-speed (240 fps) video projector (Supplementary Fig. 4). Because the presented virtual motion parallax cue automatically takes into account the rodent’s distance from the arena’s walls, virtual objects can be made to appear both inside and outside the arena’s boundaries (Supplementary Movie 1).

**Figure 1.**
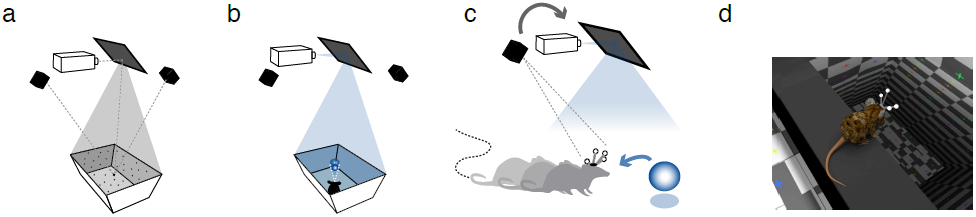
Schematic of the ratCAVE VR components. (**a**) Projector-arena mapping scheme. A dot pattern is projected onto the arena’s surface, scanned using a multi-camera 3D tracking system. A digital model of the arena is then fitted to a 3D point cloud. The projector’s position with respect to arena and tracking system is calibrated in a similar manner. (**b**) The VE projection on the arena surface, drawn from the perspective of the current position of the rodent's head, is then front-projected via the mirror onto the arena surface using a high-speed (240 fps) projector. (**c**) The 3D position a freely-moving rodent’s head is tracked by means of head-mounted array of retro-reflective spheres using a multi-camera 3D tracking system, which is used as a feedback signal for continuously updating the VE projection from the rodent’s perspective (gray arrow). By rendering the virtual environment in a 360o arc about the rodent’s head at a high frame rate (240 fps) and warping the image to match the arena’s shape, the rodent is given the illusion of a fully 3D virtual space (blue arrow). (**d**) A close-up rendering of a mouse's perspective inside a virtual environment, overlooking a virtual cliff.

**Figure 2.**
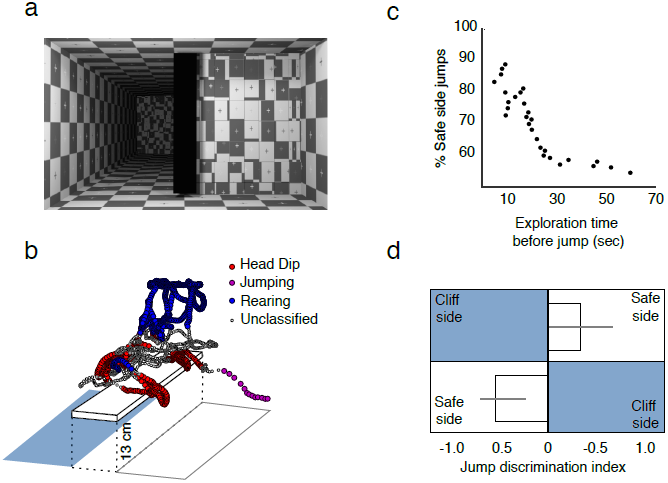
Virtual cliff avoidance experiment. (**a**) Virtual environment for the cliff avoidance test. The board was suspended 13 cm over the arena floor in the center of the arena. One half of the arena emulated a virtual cliff, a virtual floor that appeared to be 1.5 m below the arena’s actual floor. (**b**) Example trajectory and segmentation of the rat’ behavior in a single session, with three behaviors indicated by color. (**c**) Cumulative probability of jumps to safe side as a function of exploration time before the jump. Shorter exploration times were associated with a safe-side preference (see Supplementary Fig. 5). (**d**) Population jump direction statistics for sessions with jump latencies less than 18 seconds, by location of the virtual cliff. Discrimination index quantifies bias to the left or right side of the arena (see Online Methods); error bars represent a 68% confidence interval of bootstrapped means (n=25, p < .05, Fisher’s two-sided exact test).

**Figure 3.**
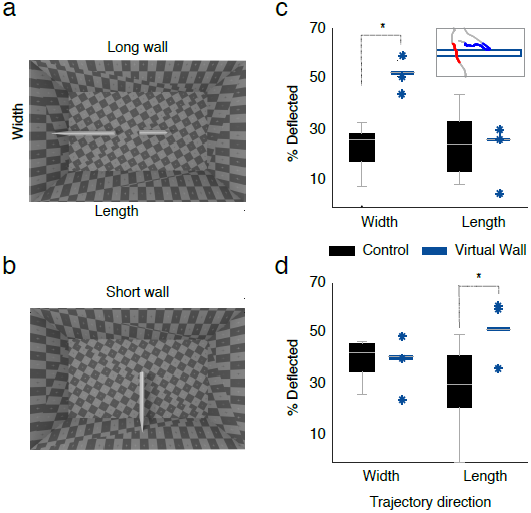
Virtual wall interaction experiment. (**a-b**) Virtual environment layout. Virtual wall in the center of arena, oriented along either the width (**a**, Long wall, first 5 min. of a session) or the length of arena (**b**, Short wall, last 5 min. of a session). (**c**) Proportion of trajectories deflected from the long virtual wall across directions of trajectories analyzed (length and width) for session conditions with the virtual wall (median, session-wise data, blue) and empty arena (control, whisker plot, black). Inset, examples of two types of trajectories: deflected trajectory (blue) and crossing trajectory (red). (**d**) Same as **c**, for virtual wall across length axis. Session-wise permutation test for significance of differences in proportions of deflected trajectories between VR (n=3 sessions) and Control (n=11 sessions) conditions in **c**: p=0.006 in width and p=0.68 in length, in **d**: p=0.65 in width and p=0.025 in length directions.

**Figure 4.**
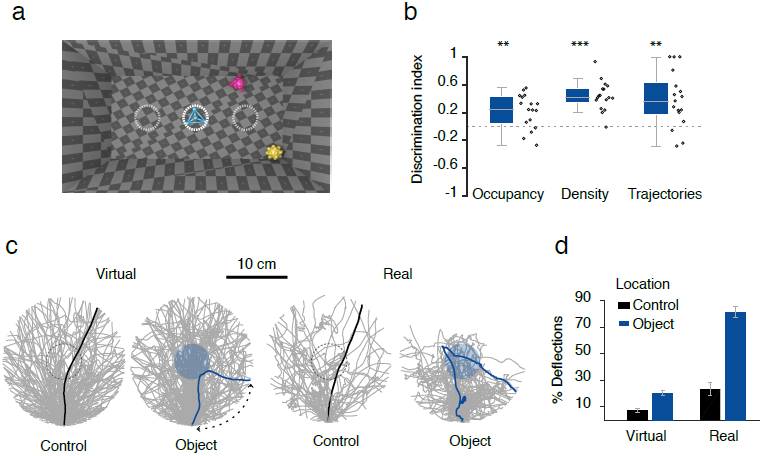
Virtual object exploration experiment. (**a**) Design of the virtual environment. Three random virtual objects were presented in a fixed relative position (“center”, “corner” and “wall”), but in a randomized arena orientation. (**b**) Object interaction analysis for the center object. Group statistics (whisker plot and individual data points, n=17 sessions) across sessions for different measures of object exploration: occupancy, occupancy density and number of trajectories converging to the center object, all expressed as a discrimination ratio between object and sham locations away from the object (dotted gray circles in a) (n=17, Wilcoxon signed-rank test, ^**^ p<.01, ^***^ p<.001). (**c**) All locomotion trajectories (gray) in the vicinity (10cm radius) of the virtual (left two plots) and real (right two plots) objects for object and sham locations. Examples of crossing (black lines) and deflection (blue lines) trajectories are shown. Trajectories have been rotated to align entry points with the bottom of the circle. Trajectories approaching within 3 cm to the object and departing at a less-than-90-degree arc from their approach vector were qualified as “deflections” (Supplementary Fig. 8d, Online Methods). (**d**) Population statistics of deflection trajectories for all conditions in c), session-wise bootstrap mean (bars) and standard deviation (error bars) of the percentage of deflected trajectories. Deflections were significantly more likely for object than sham locations exploration in both virtual (n-VRobj=72, n-sham=41, p< .02) and real sessions (n-obj=22, n-sham=16, p< .005). See Online Methods and Supplementary Fig. 8d for more information.

### Flexible design, calibration and mobility of the VR arena

Automatic arena-projector calibration ensured that the image was correctly projected onto the arena’s surface. Calibration was realized via a point cloud-modeling procedure by projecting a random dot pattern onto the arena’s surface, measuring the 3D position of each dot via a 3D tracking system, and fitting a 3D digital model of the arena to this point cloud data (Fig. 1a). This scanning process provides the flexibility to layer a VE over an arbitrary arena surface, including smooth objects inside the arena. The position of the arena with respect to the projector was continuously tracked using a set of retro-reflective spheres mounted on the arena itself, allowing the arena to be arbitrary translated and rotated during an experimental session while preserving the correct projection.

### Low latency motor-visual feedback of the ratCAVE system

Motion-to-photon (end-to-end) latency in our system cumulatively included input lag of the tracking system, the processing lag of the tracking and ratCAVE software, as well as “display lag”, the time it takes for the rendered image to be projected. Selecting fast tracking and display hardware and optimized software allowed us to achieve a motion-to-photon latency approaching 15 msec (Supplementary Fig.1 a-c). This latency is significantly lower that that of any fmVR/CAVE systems reported to date that we are aware of and additionally supplies a smoother motion stimulus than in those with lower-framerate displays (typically 60 Hz)^31,33^. Since rats rarely reached speeds of 50 cm/s during spontaneous exploration of the arena (Supplementary Fig. 1d), we expect that they were experiencing minimal, if any, latency-related cross-sensory conflicts in our system.

### Visual cues enhancing VR immersion

A large number of conflicting visual cues can exist in CAVE systems that can distract from VR immersion, which we’ve taken additional steps to decrease. First, we implemented online radiosity compensation, which equalizes the image brightness across the entire arena to decrease the visual perception of the arena itself. Second, we implemented antialiasing to decrease the perception of the individual pixels. Third, the location of the virtual light source was programmed to match the position of the projector, giving the projector the impression of simply illuminating the virtual objects, rather than creating them. Finally, to provide a richer visual scene and additional visual depth cues to the observer^34^, we implemented both diffuse and “glossy” specular reflections off the virtual objects’ surfaces using the Phong reflection model, as well as casting shadows on themselves and other objects. Additions of these visual features gave rise to a smooth and perceptually realistic VE (Supplementary Fig. 2d).

### Testing spontaneous behavior of rats in the ratCAVE

We designed a set of behavioral experiments that were aimed to explore and evaluate the degree of rats’ immersion and interaction with the VE provided by ratCAVE. In each experiment, behavior of freely-moving rats (n=3) was tested in distinct VEs that were designed to evaluate specific aspects of behavioral interaction with purely virtual elements: virtual cliff avoidance, virtual object exploration, and interaction with the virtual wall. These tasks were specifically chosen to require no pre-training or reinforcement and rely on spontaneous behavior of rodents. Benefiting from high spatial resolution tracking of position and orientation of the rats’ head, each rat’s natural behavior during each task was classified into walking, immobility and rearing based on speed and head-height features (Supplementary Fig. 6a). The three experiments were performed repeatedly across animals over several days.

### Virtual cliff avoidance experiment

Visual cliff avoidance paradigm is a classical test of visual depth perception and relies on innate behavior of the animals^35^. We designed a virtual version of this task that tests if rats avoid jumping from the virtual cliff emulated in the VE. In each 30-second session, rats were placed onto a board suspended above the arena’s floor, bisecting the arena into randomly-assigned safe and cliff sides, in which the virtual floor was at and 1.5 meters below the floor level, respectively (Fig. 2a; Supplementary Movie 3). We observed several well-defined behaviors in this task: wall-supported rearing, visual exploration of the ledges (head dipping), and the jump off the ledge towards one of the virtual floors (Fig. 2b, Supplementary Fig. 5a-c). Interestingly, rats had preference to jump to the safe side if they made their decision after short (~<20 sec exploration), but decreased this preference to chance level if longer exploration times were included (Fig. 2c, Supplementary Fig. 5d). When excluding outlier sessions (see Online Methods), we found that rats showed a preference toward the safe side regardless of the position of the virtual cliff (Fig. 2d). Thus, when exposed to the VR for a limited time rats tend to avoid it similar to real cliff avoidance paradigms.

**Figure 5.**
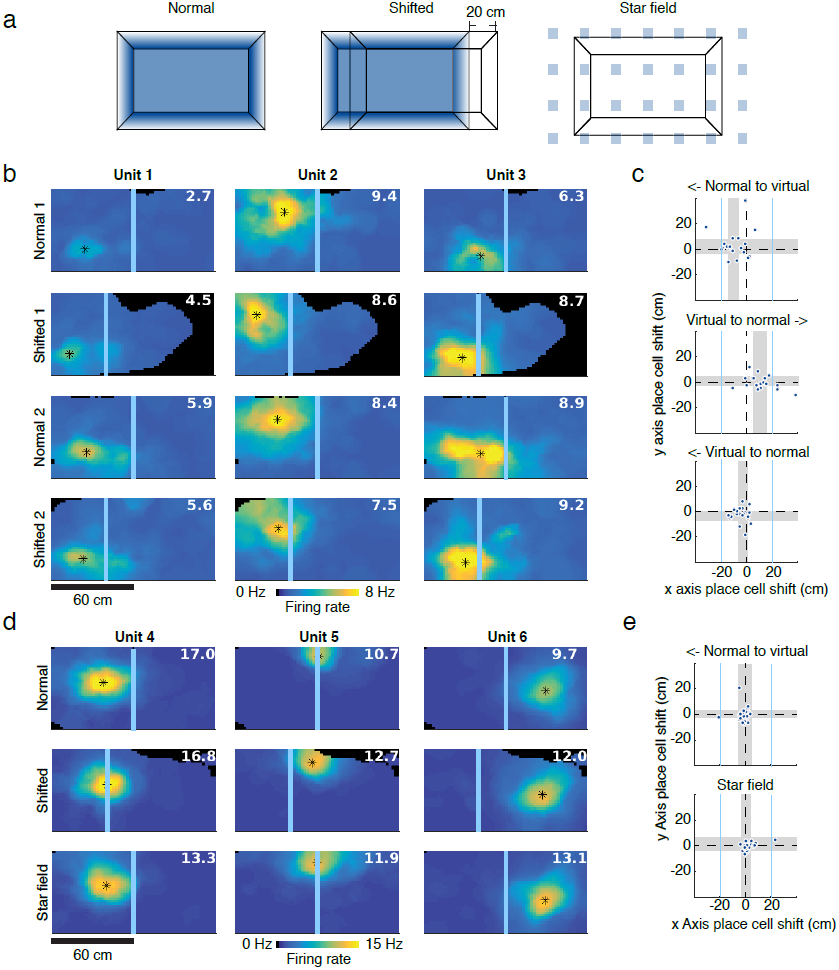
Impact of VR environment on the hippocampal spatial representation. (**a**) Schematics of the virtual environments used in the experiment. Normal (left) physical boundaries of the arena and virtual boundaries of the VE are aligned; Shifted (center) VE is displaced leftwards by 20 cm; Star Field (right), no virtual boundaries, 3D array of white virtual cubes expanding beyond arena walls. (**b**) Individual examples of place fields of hippocampal pyramidal cells showing the center position of the field (asterisk) across four sequential conditions: Normal 1 – Shifted 1 -Normal 2 – Shifted 2. The center of the virtual arena is shown as blue bars for reference. White numbers indicate the peak firing rate of the cell (spikes/s). (**c**) Analysis of the place field center shift between conditions. Scatterplots showing X-and Y-axis shift of the location of center place fields across conditions: Normal 1 to Shifted 1 (top), Shifted 1 to Normal 2 (middle), Normal 2 to Shifted 2 (bottom). Gray shadows, 95% confidence interval of population shift estimate (n=20). Non-overlap of gray bar with dotted line indicates a significant place field shift in the given axis. (**d**) Examples, as in b, for different units recorded three days later across conditions: Normal-Shifted-Star field. e) Same as c for conditions in d. Shift of place field centers between Normal and Shifted conditions was significant for length, but not width directions (Kruskall-Wallis n= 20, H=35.40, p <.001 and H=5.92, p=.21, length and width, correspondingly) and between consecutive session shifts were tested with post hoc Wilcoxon paired-rank test (n=20, in c: Normal 1 to Shift 1, W=12, p < .01; Shift 1 to Normal 2, W=15, p < .01; Normal 2 to Shift 2, W=56, p=.11; in d: Normal to Shifted, W=90, p=.67; Shifted to Star field, W=97, p=.77).

### Interaction with virtual walls

Virtual boundaries are the main elements of the VE that inform animals about topology of the virtual space^36^. In rVR systems, rats are traditionally operantly conditioned to respect the boundaries by freezing the VE upon collision of the animal’s virtual trajectory with the wall^16,21,37^. In order to investigate how naive rats spontaneously interact with virtual boundaries, we introduced a virtual wall in the middle of the arena (Fig. 3a,b). During 10-minute sessions rats were let to explore the environment. Rats displayed noticeable change of their behavior in the vicinity of the walls, as demonstrated by increased occupancy and rearing events around the wall (Supplementary Fig. 6b; Supplementary Movie 4). Interestingly, orientations of the locomotion trajectories in the vicinity of the virtual wall concentrated around perpendicular and parallel orientations to the wall (Supplementary Fig. 7b-c), indicating that the rat moved either along or towards/away to the virtual wall. Clustering of parallel orientation of trajectories near the virtual wall was similar to that in near the real wall, but was not present in the matching location in the control sessions with an empty arena (Supplementary Fig. 7). This behavior is consistent with thigmotaxis along both virtual and real walls. We further tested whether rats treated the virtual wall as an obstacle when approaching it. Locomotion trajectories approaching the virtual wall were more likely to turn away from (a “deflection” trajectory) than to cross through the virtual wall to the other side of the arena (a “crossing” trajectory, Fig. 3c), compared to the same arena locations in control sessions with empty arena, but not in the direction parallel to the virtual wall under either condition (X^2^ =48.48, n=797 trajectories, p < .001, Fig. 3c-d). Thus, rats’ interactive behavior towards the virtual wall is consistent with them responding to it as a wall.

### Exploration of virtual objects

Spontaneous exploration of the objects is the cornerstone for multitude of behavioral paradigms aimed to study perception and memory^38^. Real objects have multimodal features and affordances, but require careful and laborious handling for repeated presentation and feature manipulation. 3D virtual objects could be arbitrarily designed, manipulated and presented to an animal automatically. While rodents can perceive 3D shapes^39^ and navigate towards reward locations marked by virtual objects in rVR^22,23^, naturalistic exploration of virtual objects cannot be properly tested with any existing methods. In series of test sessions we investigated how rats spontaneously interact with the virtual 3D objects (Supplementary Fig. 8a) pseudo-randomly positioned inside the arena (Fig 4a; Supplementary Movie 5). Rats spent more time in the vicinity (<15 cm) of the virtual objects, especially in the center of the arena, with their trajectories precisely approaching the object, as compared to sham locations (Fig. 4a-b). We further quantified how rats interacted with the virtual objects on their direct approach trajectories (<10cm from the virtual object). Similar to the interactions with the virtual walls, rats’ trajectories often “deflected” from the virtual objects, reflecting that rats changed their direction of running (<90deg arc) after reaching the virtual object's boundaries (Fig. 4c-d, Supplementary Fig. 8c). Deflective nature of interaction with virtual objects was qualitatively reminiscent to that with real objects (Fig. 4c), and while less frequent, deflections were occurring significantly more often around objects than in sham locations (Fig. 4d). Rats occasionally displayed rearing and head-scanning behavior in the vicinity of the virtual objects (data not shown). Interestingly, in a fraction of sessions in which the exploration of an empty arena followed the object trial, rats showed a tendency to spend more time in the location of previous encounter with the virtual objects (Supplementary Fig. 8e).

### Effect of virtual environment on hippocampal spatial map

While, as we’ve shown above, animals immersed into the VE interact with it less reliably than with real environment and thus behavioral readout is only partially reflecting animal’s perception of the VE, internal hippocampal representation of the virtual space could provide an insight into animal’s perception of the VE^23^. Hippocampal spatial representation is believed to be anchored to multiple frames of reference, which are concurrently controlled by visual geometrical features of the boundaries and landmarks, other external sensory and idiothetic inputs, but due to physical limitation of the real environment, dissociation of the contribution of these different reference frames is difficult, and was so far mainly limited to rotations around a symmetry axis^40^. Here we illustrate an application of the ratCAVE to study complete dissociation of visual and all other multisensory systems on hippocampal spatial representation by linearly translating visual boundaries with respect to the physical environment. In the pilot experiment we recorded population of pyramidal cells in CA1/2 regions of the hippocampus (166 and 154 from two days analyzed) in a rat spontaneously exploring the arena through series of sessions in which VE was either aligned or laterally shifted by 20 cm with respect to the physical boundaries of the arena (Normal vs Shifted, Fig. 5a). Similar to the virtual wall interaction experiment, the rat interacted with the virtual boundary that appeared inside the arena in the Shifted condition at least during the first Shift session. Interestingly, population of place cells (n=20, see Online Methods for selection criteria) remapped their place fields within the arena between Normal and Shifted sessions in the direction of the VE shift (Fig. 5b-c). The effect decreased over consecutive alternating sessions and following multiple exposures to the shifted VE (3 days later) place cells showed no remapping between Shifted and Normal conditions (Fig. 5d-e). We tested if any visual information associated with VE boundaries is contributing to the stabilized spatial map by immersing the rat into the VE that was unrelated to and expanded beyond the physical boundaries of the arena. This VE as well had no effect on the place field position (Fig. 5d-e, bottom). Thus ratCAVE is sufficiently immersive to enable visual input control of hippocampal spatial representation, but progressive exposure to the conflict between visual and other multisensory inputs enabled by ratCAVE can result in complete independence of the hippocampal spatial representation from the visual input^41,42^.

## DISCUSSION

We presented a ratCAVE system for freely-moving rodents that builds on and extends previous developments of fmVR systems in arthropods and fish^28-31^ to provide a high-performance general cognitive science VR research platform by implementing a combination of methods that provide realistic visual environments, low-latency and high-precision closed loop feedback to animals’ head, and flexibility of the shape and mobility of the arena. Using more complex lighting models, including diffuse and specular reflections and self-shadowing, provides new visuo-spatial cues for virtual environments and increases immersion^34,43^. In humans, sensory conflicts resulting from out-of-phase feedback to rapid head motion arise when motion-to-photon latency of the VR system is larger than ca. 50 msec, resulting in decreased performance in spatial navigation, spatial perception, and sense of self-motion in the VE^32^; to counter this effect, we've implemented a low-latency visual update loop (240 fps, 15msec “motion-to-photon” lag) to decrease mismatch between vestibular, proprioceptive, and visual self-motion cues, essential for proper self-motion detection and functioning of the head-direction system^44,45^.

There are pressing improvements needed to further increase immersion in VR systems used in neuroscience research. While rVR immersion requires animals to ignore lacking or mismatching sensory inputs, immersion in fmVR is associated with the minimal conflict between visual and other senses. However, both rVR and fmVR systems suffer from the cross-sensory conflict upon collision of animal’s trajectory with the virtual boundary and can break immersion. In rVR setups, the solution has been to simply stop visual update while still allowing rodent locomotion, creating a locomotion-visual mismatch upon impact^16,23^. In fmVR, a similar mismatch occurs when the virtual and real surfaces are not matched and are directly sampled by the animal. Such situations require a careful selection of virtual environment, arena design and method to match the research questions at hand. A few improvements can be considered in the ratCAVE. First, VE objects and boundaries can be made inaccessible to the animal by projecting them outside arena walls or across the gap. Second, ratCAVE calibration procedure allows for projecting virtual objects on smooth shapes inside the arena, thus aligning them with real countrerparts, enhancing VR immersion via all three avenues: naturalistic interaction (via touch and smell), increased cue salience, and reducing cross-sensory mismatch upon virtual object contact. Third, electrical or optogenetic stimulation of olfactory or somatosensory system^46,47^ can be used to provide congruent multisensory feedback. Similarly, use of visuo-acoustic VR can be provide more cohesive VE^22,48^. In addition to motion-dependent monocular depth cues, static binocular depth cues based on stereoscopy are also important for forming an accurate 3D space percept^43^, a point currently ignored in rodent VR studies. Thanks to precise head-based projection, the ratCAVE system can be extended to generate stereo VE via implementation of head-mounted shutter glasses to provide alternating images to the left and right eyes of the exploring rodent. Many of these improvements can be added onto existing rVR and fmVR systems to increase VR immersion in those setups. Further integration and cross-insemination of open-source fmVR and rVR developments in diverse animal models will enable a broad spectrum of neuroscientists to use these systems.

Freely-moving virtual reality represents an improvement in VE immersion over rVR, considered as an enhancement of naturalistic interaction mechanisms with the virtual environment, an increased salience of sensory cues associated with the virtual environment, and a minimization of cross-sensory conflict. Naturalistic interaction with the virtual environment is enhanced in fmVR by simply allowing the full range of movement in an unmodified space, without training or postural alteration, while in rVR, locomotion and virtual object interaction must be simulated via running on a spherical treadmill. Self-motion cues through the virtual environment are enhanced in fmVR by providing higher-frequency and shorter-latency feedback to head motions in the virtual environment alongside the lower-frequency locomotion behaviors, while rVR only provides locomotion feedback. In contrast to rVR that assumes a stationary head in the virtual projection, fmVR system minimizes cross-sensory conflict by providing feedback to head motions, as well as by matching changes in olfactory, tactile, and auditory real-world inputs to self-motion in the virtual world. Finally, fmVR systems do not require operant training and habituation procedures used in rVR systems.

We demonstrated that a ratCAVE VR system for freely moving animals can be successfully applied to a number of behavioral paradigms not possible with conventional rVR systems. Untrained rats freely behaved and spontaneously interacted with virtual environment by approaching, exploring and leaving virtual objects and walls, displaying thigmotaxis along virtual walls and avoiding a virtual cliff. We further used ratCAVE system to illustrate how contribution of the virtual visual input to hippocampal spatial representation can be strong upon first exposure to VE mismatched with the physical world, but becomes negligible after repeated exposure of the rat to cross-sensory conflict. These experiments and design features of ratCAVE described above pave the way to a large body of future applications.

First, high spatio-temporal resolution of 3D tracking of the rodent’s head, which can be extended to include the full body, enables quantitative analysis of the natural behaviors of the rodent during VE exploration, which significantly extends level of analysis possible using two-dimensional locomotion information provided by conventional tracking in 2D space or the treadmill measurement in rVR. Second, ratCAVE's “trackable” arena also enables vestibular perturbations during VR experiments via arena movement, enabling studies on vestibular system function and visuo-vestibular binding in behaving rodents. Third, fmVR's ability to incorporate a three-dimensional element into operant conditioning tasks increases the range of motor affordances of digitally-rendered learning stimuli, which have their own benefits of flexibility and timing control^49^. Integrating these improvements into VR setups will enable new methods in research areas such as learning and memory, perceptual decision-making, and 3D-rotation and object perception^39^. Fourth, the automated nature of head tracking allowing for online behavior analysis, operant conditioning, and fmVR enable high-throughput and automatic behavioral testing in a colony of animals^50^ across a large variety of tasks, such as perceptual, incidental and motor learning, spatial memory paradigms, to name a few. Importantly, use of automated fmVR behavioral paradigms allows their standardization, reproducibility of results independent of experimenters or setup. Finally, combined with neural recording and manipulation ratCAVE enables the detailed investigation of the mechanisms of spatial coding. Manipulation of the arena boundaries provides a powerful tool to study for multisensory nature, remapping and attractor properties of the spatial representation^51^.

Low latency, unmatched by any other system for freely moving subjects, and rich visual features make ratCAVE appealing for use in human subjects. Translation of experimental paradigms and physiological validation of psychophysical experiments from humans to animals and back could enable validation and further development of diagnostic and rehabilitation procedures for the vestibular or neurodegenerative disorders in animal models^52,53^. ratCAVE opens new ways to study sensory-motor systems in their natural dynamics while having flexibility in manipulating the sensory feedback not possible in real life.

## ACKNOWLEDGMENTS

We thank Esmi Zajaczkowski for help in experiments and design of the 3D virtual objects, Nicolas Kuske for help with programming API for Optitrack camera control. This work was supported by the Deutsche Forschungsgemeinschaft (DFG) via the Werner Reichardt Centre for Integrative Neuroscience (CIN, Tuebingen), Munich Cluster for Systems Neurology (SyNergy, Munich), Priority Programs 1665 and 1392 and by the German Ministry for Education and Research (BMBF) via Grant Number 01GQ0440 (Bernstein Center for Computational Neuroscience Munich).

## AUTHOR CONTRIBUTIONS

N.A.D.G. designed and implemented the ratCAVE system, designed and performed behavioral experiments in the VR; J.J.G. and E.B.H. performed electrophysiological experiments recordings in the VR and behavioral experiments with real objects exploration; all authors contributed to data analysis; N.A.D.G., E.B.H and A.S. wrote the paper. All authors discussed the results and commented on the manuscript.

## ONLINE METHODS

### ratCAVE VR system

**Hardware setup.** Our setup consisted of a rectangular arena with dimensions 115cm × 65cm (L, W) and walls 40cm high, angled at 70 degrees to increase the projected image's surface area and brightness. A set of 12 cameras (OptiTrack, NaturalPoint Inc. U.S) was used to record the 3D position of retro-reflective spheres, six Prime 17W (360 fps) and six Prime 13W (240 fps). A projector with 240 fps frame rate (VPixx Technologies Inc., Saint-Bruno, Canada) was mounted to the ceiling. An optically-flat aluminum-foil projection mirror (100cm × 75 cm, Screen-Tech), slanted 45 degrees, was suspended from the ceiling on an adjustable frame for accurately fitting the projected image onto the whole surface of the arena. This setup was installed inside an isolating acoustic chamber (Supplementary Fig. 4).

**Software.** The ratCAVE VR system depends on many pieces of software to work; interactions between each software component are diagrammed in Supplementary Figure 3. Virtual environments are modeled and exported to file in a 3D modeling program, Blender 3D (Supplementary Fig. 3a). Coregistration of the arena and projector with the tracking coordinate system is performed via a custom Python command-line program package called “ratcave_calibration”, which uses a custom Python API called “MotivePy” to access and controlling our Optitrack camera array while using a custom Python 3D graphics utility package called “Fruitloop” to render the point cloud from the projector (Supplementary Fig. 3, “Grey Zone”). Fruitloop provides a user-friendly interface for modern OpenGL rendering techniques, and its “Get Data, Update Camera, Render VE” event loop forms the core engine of a ratCAVE virtual reality session. Cubemapping, lighting, and antialiasing are done via OpenGL FrameBuffer objects and shader scripts supplied with Fruitloop. VR Experiment scripts are written in Python, using a custom network client called “NatNetClient” to obtain Optitrack camera data in real-time and Fruitloop to render the virtual scene (Supplementary Fig. 3, “Blue Zone”). Because all software used in the ratCAVE VR setup is comprised of loosely-connected specialized parts, the software developed by the lab is generalizable to a variety of different setups, enabling other labs to substitute like-components to build a VR setup that matches their hardware.

**Code availability.** All code used in implementation of the ratCAVE is freely available for use and modification via Github and installable via the Python Package Repository. The Fruitloop package can be found at https://github.com/neuroneuro15/fruitloop, MotivePy at https://github.com/neuroneuro15/motivepy, NatNetClient at https://github.com/neuroneuro15/natnetclient, and ratcave_calibrate at https://github.com/neuroneuro15/ratcave_calibrate. Associated documentation and usage tutorials are available at fruitloop.readthedocs.io and in Supplementary Documentation. All custom-written code used for analysis of results presented in this study are available from the corresponding author on reasonable request.

**Data availability**. The datasets generated during and/or analyzed during the current study are available from the corresponding author on reasonable request.

**VR implementation.** We tracked the rodents' head position and orientation by imaging a head-mounted, 3D-printed plastic skeleton of four retro-reflective spheres (6-8mm in diameter). Commercial software (Motive, NaturalPoint Inc., USA) isolated these spheres' positions in each camera's imaging data and reconstructed the 3D position and orientation of the rigid body (Supplementary Fig. 3, “3D Tracking Software”). Rodent head position was then logged for offline analysis and sent over the network to the VR system's experiment script via a custom python package (NatNetClient) for visual stimulus update (Supplementary Fig. 3, “Python Optitrack Client”). The ratCAVE VR engine Fruitloop receives the current position of the rat’s head from NatNetClient, updating the virtual scene from the rat’s perspective, generates the projected image using a cube-mapping algorithm (Supplementary Fig. 2a-c), performs per-fragment lighting calculations (Supplementary Fig. 2d), and antialiases the resultant video output via custom OpenGL shaders (Supplementary Fig. 3). The resultant image is then projected onto the arena via the video projector.

**Latency measurement.** Motion-to-photon latency was explicitly measured using the following setup ^54^. A reference point, representing a VR observer, formed by a set of three retro-reflecting markers and a small LED, were attached to a bar that was rotated in the horizontal plain around a fixed point inside an arena by an AC motor and was tracked as described above. The VR system was programmed to generate a white spot that was offset in the horizontal plain from the reference point that would follow a reference marker. VR spot was thus rotating in the horizontal plain following the rotation of the reference LED point. Both LED and VR spots were imaged using high-speed-camera (Prime, Photometrics) at 250 Hz. The image stack was processed to detect both spots (Supplementary Fig. 1a) and temporal trajectories of X and Y coordinates of both reference and VR spots, which were analyzed to detect temporal offset between them using cross-correlation function (Supplementary Fig. 1b). The angular speed of rotation was varied between trials, and the resulting linear speed (tangential) was computed and used for latency-speed analysis (Supplementary Fig.1c).

### Animal experiments methods

All procedures complied with the European Communities Council Directive 2010/63/EC and the German Law for Protection of Animals and were approved by the local authorities, following appropriate ethics review.

**Subjects.** Three 6-month-old male Long-Evans rats (Charles-River, Germany) were used for the analysis of spontaneous exploratory behavior in virtual environments, and three rats were used for analysis of spontaneous exploration of real-world objects. An additional rat was used to record hippocampal neural activity in a virtual environment, as described in the “VE Shift Experiment” section. All rats were allowed *ad libitum* access to water and food. All rats were extensively handled by the experimenter prior to behavioral experiments in order to minimize stress.

### Behavioral experiments

We recorded the spontaneous behaviors of three rats in three virtual environments. Each session, conducted twice per day over one week, consisted of two phases: a one-minute visual cliff session and a ten-minute arena exploration session, between which the rat was removed from the arena. Same-day sessions were separated by a minimum of 5 hours; the first in the middle of the rat's light cycle, and the second at the beginning of its dark cycle (labeled in Supp. Figure 5 as “Midday” and “Evening” sessions, respectively). Arena exploration sessions could contain either virtual objects for exploration, virtual walls, or no virtual objects at all (control condition; the projector remained on and a stationary checkerboard pattern remained projected on the arena).

**Virtual cliff experiment.** In each virtual cliff session, the experimenter placed the rat on a 14 cm-wide board suspended 13 cm above the arena’s floor, bisecting the arena into two halves. Two virtual floors were randomly assigned and projected onto the arena halves, either 1.5 meters below the actual floor level (the cliff side) or at the same level as the floor (the safe side). We enhanced the motion parallax stimulus by adding a slight height variation to the floor texture; this also helped reduce the chance of side selection by the presence of virtual motion. After some visual exploration of their environments, rats jumped down from the suspended board to the arena floor on one of two sides and were allowed to explore for 15 seconds, after which the experimenter removed the rat from the arena. Each session lasted a maximum of 90 seconds. Cliff avoidance behavior was interpreted by observation of the rat jumping down from the board on the safe side, after a period of visual exploration of the arena, with jump side observation counts calculated as a discrimination index {(safe-cliff)/(safe + cliff)}.

**Virtual wall experiment.** During the virtual wall sessions, rats were allowed to freely explore the arena for 10 minutes. A virtual wall extended from the center of the arena, dividing it across its length (short wall) or its width (long wall). Each rat was exposed to both walls for five minutes (long followed by short wall) in a single session.

**Virtual object exploration.** During object exploration, rats were allowed to freely explore three different virtual objects, each roughly 6 cm in diameter and randomly selected from a pool of 11 custom-designed 3D models (Supplementary Fig. 8a). The objects were placed in the nodes of triangular configuration “corner”, “wall”, and “center” (Fig. 4a), which was pseudo-randomly rotated between trials. In some of these sessions, the objects displayed either a shrinking, rotating, jumping, or running animation when the rat came within 15 centimeters of the object’s center, with the goal of increasing rodent engagement with the objects, although this factor was ignored in the analyses due to low sample size.

**Virtual environment shift experiment.** During the VE shift experiments, the rat was allowed to explore the arena for ~10 minutes. In consecutive sessions, the rat experienced two conditions: a “Normal” condition, where the arena had virtual walls (checkerboard pattern) matching entirely in space with the real walls, and a “Shifted” condition, where the VE was shifted 20 cm along the arena's length. This effectively resulted in the shift of one virtual wall to the outside of the arena and another to the inside of the arena, similar to the virtual wall experiment. The rat was exposed to the arena and these two conditions twice for the first time on Day 1 (Normal1, Shift1, Normal2, Shift2 in Fig. 5 b-c) and then repeatedly to the same conditions, as well as other VR manipulations in several sessions on Days 2 and 3 (data not shown). On Day 4 two sessions were recorded under Normal and Shifted conditions and an extra session under condition “Star field” was introduced (Normal, Shift and Start field in Fig. 5d-e). This condition consisted a 3D grid of repeating white cubes, which extended 1 meter beyond the walls and floor of the arena.

**Surgery and electrophysiological recordings.** Rats were anesthetized with a three-component mixture (Fentanyl .005mg/kg, Midazolam 2mg/kg, Medetomidine .15mg/kg); this compound also provided analgesia for the first part of the procedure. A 1.5% concentration of isoflurane in oxygen was used to maintain depth of anesthesia for the rest of the surgery. In animals used for behavioral assays, a small screw was fixed into the skull to provide support for our head post. In one rat, a silicon probe (NeuroNexus, Buzsaki 32 design, 4 shanks, 8 sites ~25um vertically spaced) was implanted following procedures described elsewhere ^55^. Briefly, a cranial window of ~2 mm^2^ was opened, centered on the following coordinates from bregma:-3.36 mm AP and +2.6 mm ML. The silicon probe, mounted on a custom-made microdrive, was inserted in the center of the craniotomy with the shanks aligned parallel to the septo-temporal axis of the hippocampus (45 degrees parasagittal). The probe was lowered to a distance 1mm from the surface, and the drive was affixed to the skull. After the rat recovered from surgery (1 week), the probe was lowered 50-150 microns daily until we observed the typical profile of activity of CA1/CA2 cell layer; namely, spiking activity and ripple oscillation signal in the LFP. Histological verification of the location of the recording electrodes was done after the conclusion of the experiments (data not shown).

**Data acquisition and processing**. Extracellular signals were amplified and filtered by multi-channel preamplifiers (Plexon, 20×, 1-5000 Hz). Wide-band extracellular and intracellular signals were digitized at a 20 kHz sampling rate with 16-bit resolution and stored for offline analysis using a multichannel acquisition system (DigiLynx, Neuralynx). Raw data were preprocessed using a custom-developed suite of programs (neurosuite.sourceforge.net). The wide-band signal was downsampled to 1.25 kHz and used as the local field potential signal. For spike detection, the wide-band signal was high-pass filtered (>0.8 kHz). Single units were isolated semi-automatically by a open-source spike-sorting program KlustaKwik (http://klustawik.sourceforge.net) ^56^ and refined manually using open-source GUI software (http://klusters.sourceforge.net; http://neuroscope.sourceforge.net) ^57^. The quality of isolated single units was confirmed by an isolation distance metric and a clean refractory period.

### Data Analysis

All data analysis was performed using custom-written code in Python and Matlab (Mathworks, Inc.).

**Data representation and statistics.** If not described in the figure legends data summary are plotted using Matlab boxplot (whisker plot) showing median, 25/75 percentile bar and whiskers extending to +/-2.7σ outliers are shown with small crosses. Non-parametric and resampling statistical analysis methods were used in all cases and are specified in figure legends and methods sections below. Due to use of non-parametric methods assumptions (e.g. normality of the distribution), associated with parametric tests were not tested. Size of experimental animal sample to ensure adequate power could not be determined prior to the study, since no parameters of analysis could be predicted *a priory*. Instead of increasing animal sample size, we repeated individual experimental sessions (virtual cliff and object exploration; virtual wall condition was not repeated due to recognized interference between VR sessions due to potential memory effect). No animals or sessions were excluded from the analysis. Randomization and blinding was not performed as all animals were subjected to all tests conditions as well as control sessions.

**Behavioral state classification.** Behavioral state of the rat was classified based on the speed and height of the head. Using data-derived thresholds for these variables, we defined running (speed > 3cm/s & height < 13.4cm), immobility (speed <= 3cm/s & height < 13.4cm) and rearing (speed <= 3cm/s & height > 15.4cm).

**Virtual cliff avoidance analysis.** Rat behavior was segmented based on head-tracking data into supported rearing on the arena walls and general exploratory behavior In addition, visual exploration was associated with head dips, which were detected as trajectories of the head extended within 1cm from the board. Jumps were detected as trajectories that depart from the board and land on the floor. A rat's landing after jumping down from the board was detected based on the height of its head (threshold < 7 cm). Example sessions' time courses are shown in Supplementary Fig. 5a. Since rats spent a variable amount of time across sessions performing supported rearing (M=24.8% of trial, SD=15.5%), likely trying to escape the arena or look outside the arena, we chose to remove these periods from the decision time estimation analysis, yielding an exploration time measure before the jump event, which we used to analyze the effect of time spent exploring the VE prior to the jump side decision behavior (Fig. 2b, Supplementary Fig. 5d). Excluding supported rearing did not make any qualitative difference in the outcome of the statistical analysis. Cliff avoidance behavior was interpreted by observation of the rat jumping down from the board on the safe side, after a period of visual exploration of the arena, with jump side observation counts calculated as a discrimination index [{(safe-cliff)/(safe + cliff)]}.

**Virtual wall experiment analysis**. Only periods of continuous locomotion were considered in the analysis of trajectories. When rats locomoted to the vicinity (within 7.5 cm) of one side of the virtual wall, the trajectory was counted as a “Crossing” if they continued through the wall and reached the threshold on the other side; if they returned to the same side of the arena where they entered the vicinity, the trajectory was counted as a “Deflection”. A chi-square test of independence on deflection and crossing counts was used to test for an increase in total number of deflection trajectories for the VR wall condition over control sessions in the same locations. To test if the proportion of deflecting trajectories exceeded chance level for each condition (Long / Short), we permuted (6000 permutations of 3-session means) the proportion of deflections for VR (n=3) and Control (n=11) sessions combined to generate a null distribution, and computed an empirical one-sided p-value for the actually-found percentage of mean for trajectories deflecting off the virtual wall for both Long and Short VR conditions.

Thigmotaxis behavior, i.e. locomotion along and near the wall, was quantified using analysis of the angle of instantaneous velocity vector with respect to the virtual wall and same axis for control conditions with no wall present. Instantaneous 2D velocity vector was smoothed (500 msec boxcar) and the angle between this vector and the wall was extracted. Joint probability density function between distance of the animal instantaneous position and angle to the wall was computed (Supplementary Fig. 7b,c). Presence of the mode at 0 radians for animals positions close to the virtual wall (blue line) and real wall (black line) indicated comparable thigmotaxis behavior. Control conditions, in contrast to virtual wall conditions lacked this mode. A second mode around pi/2 for distances spanning large range from the virtual wall indicates trajectories crossing and deflecting from the virtual wall, as analyzed and represented in Figure 3.

**Virtual object exploration analysis.** Object exploration was quantified using a set of metrics aimed to measure rats’ exploration of the virtual objects’ locations. We used progressively more refined measures to quantify animals' exploration of the virtual object. First, an *occupancy* of the object vicinity, i.e. probability that rat is located within 15 cm from the object, was used as a crude measure to assess the general preference of the animal to be near the virtual objects. Second, the *occupancy density* at the object location, computed as a ratio of occupancy within 5 cm to that within 15cm of the object’s center, was used to measure the selective localization of increased occupancy within the direct vicinity of the object. Third, we analyzed the proportion of *trajectories* that entered the vicinity of the object (10cm radius) that reached within 3cm of the virtual object’ center. To control for the significance of this effect against random locomotor activity, which is naturally constrained and interacts with the arena walls, we first considered using control sessions that contained no objects within the arena. Surprisingly, we found an increased occupancy at virtual object locations compared to the rest of the arena in these sessions (Supplementary Fig. 8e), potentially reflecting a memory effect of the animals for the location of the objects. To avoid these inter-session interactions, all further trajectory analyses weredone against a within-session control “sham” location, paired to each virtual object on the opposite side of the arena (Fig. 4a; Supplementary Fig. 8b, top). For all measures of object exploration we constructed a discrimination index [DI = (VR – Sham) / (VR + Sham)] and tested using a Wilcoxon signed-rank test for significant differences from zero. Consistent with observations in both other studies utilizing real-object discrimination tasks and our own analysis (Supplementary Fig. 7), animal behavior in the vicinity of both the arena and virtual wall boundaries was heavily biased to thigmotaxis. In addition, we observed a high rate of supported rearing next to the walls and, especially, in the corners (Supplementary Fig. 6b). These factors heavily contaminated and made insensitive most measures of spontaneous exploration of the objects located next to the wall and in the corner. Consistently, we found that occupancy times for the object vs sham were significantly higher for the center object (Z=2.70, p<.01, Fig. 4b), but not the wall object (Z=1.42, p=.08, data not shown) nor the corner object (Z=-0.52, p=.70, data not shown). Occupancy density was significantly different from sham for the center object (Z=3.55, p<.001, Fig. 4b) and corner object (Z=2.13, p<.05, data not shown), but not the wall object (Z=0.56, p=.29, data not shown). Locomotion trajectories approaching the object (within 10 cm) were also more likely to pass through the VR objects than their sham pairs for the center object (Z=2.889, p<.01, Fig. 4b) and wall object (Z=2.130, p<.05, Supplementary Figure 8b,c), but not the corner object (Z=1.008, p=.15, Supplementary Figure 8b,c).

The rats sometimes interacted with the virtual objects and then changed their running direction. To quantify this behavior, we introduced a notion of trajectory “deflections” from the object (see Fig. 4c for trajectories examples). We analyzed the relationship of the arc angle made by trajectories entering and leaving the 10 cm circle around the object, a “deflection angle”, with the shortest distance between the trajectory to the object. If a trajectory approached the object closely and its deflection angle was acute (< 90 degrees), we qualified it as a “deflecting”, while obtuse (> 90 degrees) deflection angles were qualified as “crossing”. As trajectories not reaching the proximity of the object are progressively associated with smaller deflection angles, we set a conservative cut-off distance of 3 cm to define a trajectory as deflecting. Thus, deflecting trajectories are those that fall in the region of less than 3 cm and less than 1.56 radians, displayed in Supplementary Fig. 8d. We compared the proportion of “deflecting” trajectories for sham and object-containing locations using an object-label-shuffling permutation test (Supplementary Fig. 8d, right column). As the arena wall blocked trajectories for the other two object positions (“wall” and “corner”), deflection trajectory analysis was only possible for the center object. To compare this newly-introduced measure of rat interaction with the virtual object with that for the real objects, we performed an identical analysis in a separate set of data from 3 rats exploring real objects in a cylindrical arena.

In the fraction of sessions where virtual objects were programmed to be interactive, i.e. displayed either a shrinking, rotating, jumping, or running animation upon the rat’s approach, we observed increased exploration in the vicinity of the objects (data not shown). As this behavior was variable across animals, our data lacked sufficient power to statistically assess this effect.

**Brain state segmentation.** Hippocampal activity was segmented into two states: theta and non-theta. An HMM Gaussian mixture model based on the hippocampal CA1 pyramidal layer spectral power ratio between the 6-12 Hz band and the sum of the 1-5 Hz and 15-18 Hz bands of the whitened LFP was used to separate theta and non-theta states. All further analysis of the hippocampal place cells was constrained to theta-associated periods.

**Place cells analysis.** Only hippocampal pyramidal cells with place fields that were active in the arena were included in the analysis. Spike width and firing rate were used to separate pyramidal cells from interneurons. In the sessions used in this paper, 309 of 367 cells were classified as putative pyramidal cells (Day 1: 166 of 168 cells; Day 4: 143 of 182 cells). Place cells were defined as putative pyramidal units with a place field peak firing rate of at least 3 Hz, having less than three spatially-separated firing rate peaks in all trials, and maintaining a stable spike waveshape across all sessions of the day. After filtering based on these selection criteria, 39 total pyramidal cells for the two days (20 and 19 cells, respectively) remained.

Place fields were calculated based on a k-nearest neighbor algorithm, which selected for periods in which the speed of the rat's head was greater than 5 cm/s and intersected with periods of theta oscillation state. The k-nearest neighbor estimate of the mean firing rate was calculated given the position of the rats head and each unit's smoothed firing rate. The unit firing rate was smoothed using a 800 ms rectangular window, convolved with the time-resolved spike histogram and downsampled to 30 Hz. The maze was binned with 2 cm square bins. For each bin, the smoothed unit firing rate was sorted by its distance to the bin center. The first 300 nearest-neighbor time bins were collected and averaged to derive the mean rate of that bin. Bins with less than 300 neighbors within a radius of 12.5cm were assigned to be empty. This procedure provides a data-adaptive and robust estimate of the spatial rate map in contrast to conventional estimation methods (ratio between spatially smoothed spike count and rat occupancy maps). Qualitatively, though, both measures gave the same results.

The procedure used for place field map estimation has additional benefits, as it allows robust estimation of the parameters of the place field based on the bootstrap procedure. The variance of the place field center was estimated by bootstrapping each unit's 1000 random subsets of two-second chunks of the rodent's trajectories (75% of total Trial time). The place field center at each iteration was calculated by thresholding the rate map by the firing rate at the 95 percentile of all iterations. All bins above the threshold were assigned a 1, and all other values were assigned a 0. The above-threshold bins were segmented using Matlab’s bwboundaries function into spatially contiguous patches, each of which represented a place field. The area, the rate-weighted center of mass, and the maximum and minimum firing rate were calculated for each patch. Only the main (largest and highest firing rate) field was used for further analysis. The location of the peak rate within the patch was computed for each bootstrap sample, and the resultant mean estimate was used as an unbiased estimate of the x-y position of the center of the place field and used for the further analysis of place field remapping. To quantify the effect of the VE shift on the place fields of the active population of place cells, we computed the displacement of the place field center between consecutive sessions (Normal to Shift, Shift to Normal etc). The Kruskall-Wallis test was used to find an overall difference in population means between sessions along each axis of the arena, and significant axes were probed for individual differences between sessions using a Wilcoxon paired-rank test with p-values corrected for multiple comparisons using Benjamini/Hochberg False Discovery Rate method.

---

**Supplementary Figure 1.**
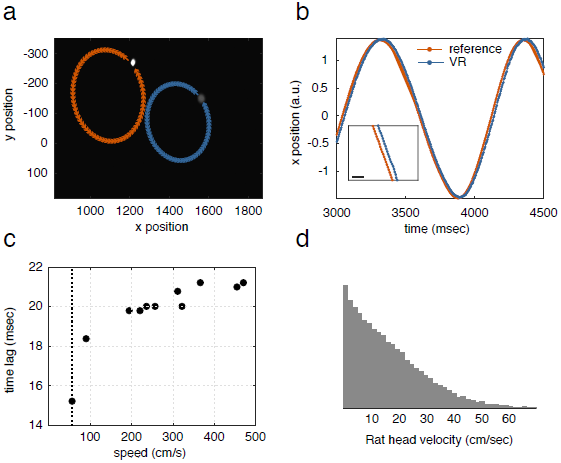
Motion-to-photon lag measurement of ratCAVE system. (**a**) Raw image of the latency data collection procedure. A tracked object (reference point, left spot, orange) and its x-axis offset VR-represented projection (virtual point, right spot, blue) were recorded using high-speed camera as the reference point was rotated about a central point at different speeds. Arrows show the direction and shape of the reference and virtual points’ trajectories. (**b**) Example of the time courses of the x-axis project of the reference (blue) and virtual (red) points. Inset, magnification of a section of time course. Note the delay between the two time courses, scale bar 10 msec. (**c**) Time lag between the reference and virtual points as a function of linear (tangential) velocity of the reference point motion. Note slow increase of time lag with speed of the reference point above 100 cm/s, with a minimum latency of 15 msecs in the range of the head velocity of rats. (**d**) Distribution of head velocity in rats, note that all movement are contained within 60cm/sec (doted line in c).

**Supplementary Figure 2.**
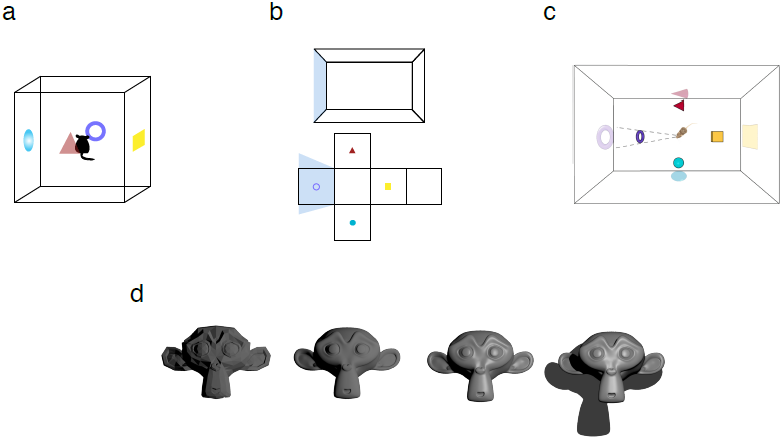
Cube-mapping and image rendering. (**a-b**) Schematic of the image-warping transformation of the rat’s perspective view of the virtual environment to the projection on the arena. The image warping algorithm involves three steps: (**a**) The virtual world (consisting, in this example, of four colored 3D objects) is rendered 360 degrees about the rat's head position on the faces of the cube using a cube-mapping algorithm, (**b**) each wall's relative position to the rat is mapped to this 3D virtual world,and (**c**) all arena surfaces (walls and floor) are then warped from the perspective of a video projector mounted above the arena. This process is repeated every frame, maintaining the VR-rodent-arena despite movement of virtual objects, the rat, or the arena itself. (**d**) 3D lighting algorithms employed by ratCAVE to increase spatial visual cues and visual richness of the virtual environment. Improvements are successively applied to the object, from left to right. First, diffuse reflections increase object brightness on parts of the object facing the light. Second, high-resolution objects are used, with smoothed surfaces, to further increase object detail. Third, specular reflections are added to provide subject-object-light triangulation cues. Finally, shadows are added to provide inter-object distance cues.

**Supplementary Figure 3.**
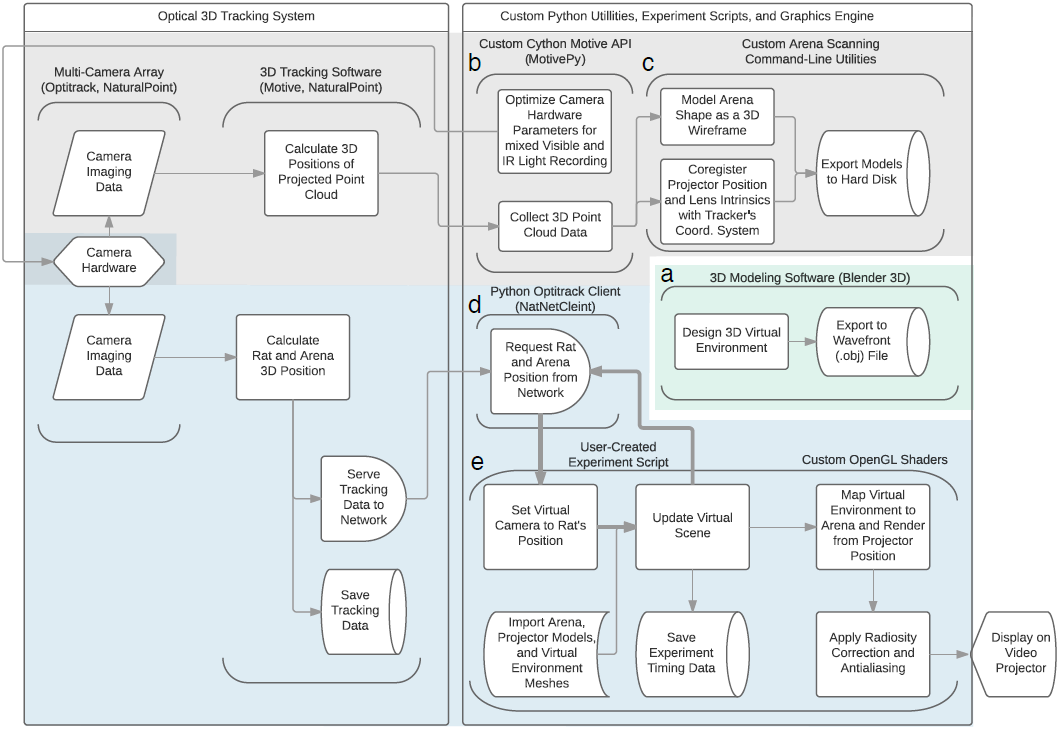
RatCAVE hardware-software components flowchart. Each component, depicted as vertical parenthesis, takes information from one source and sends information to another source; information flow is depicted in direction of arrows. Detailed operations of each component are depicted as blocks, and software components are labeled by letter. (**a**) Blender 3D. The virtual environment is created before the experiment using 3D modeling software (right-center module) for loading into the VR experiment script. (**Gray Zone**) Tracking and Setup Coregistration. A Multi-camera array sends imaging data of the rodent's position on each camera to 3D tracking software, which combines the data from each camera image into a single 3D location and sends that position to the ratCAVE environment (left, “Optical 3D Tracking System”). (**b**) MotivePy. The cameras' settings can be modified directly in a Python environment to make visible-light collection possible, a necessary step for arena scanning and projector calibration. (**c**) ratcave_calibrate. Two command-line programs are used for arena scanning and projector calibration. The arena scanning program projects a moving grid of white dots on the arena surface, collects the 3D positions of the projected points via the camera array, and fits the resultant point cloud to a 3D mesh model of the arena. The projector calibration program maps single points displayed from the projector onto the 3D position of the arena, one at a time. It then uses OpenCV's camera_calibrate tool to use these mappings to find the position of the projector in the camera array's coordinate space. (**Blue Zone**) VR Engine. (**d**) NatNetClient. Rat position data is collected in real-time from the camera array and brought into the Python environment, for use in VR experiment scripts. (**e**) Fruitloop. The virtual environment (VE) is rendered in a Python 3D graphics engine. The VE is loaded from file (created in Blender 3D), and on each display frame, using the rodent position data to move a virtual camera to the rodent's position in a virtual environment (blue zone). This process, encompassing the core of the VR engine, (get rodent position, move camera, update and render scene) occurs in a loop, repeated each frame, with the frames themselves sent to the GPU for arena mapping and shading (examples on Supp. Figure 1d) and then to the video projector (bottom-right corner). See the “Software” section in Online Methods for more details.

**Supplementary Figure 4.**
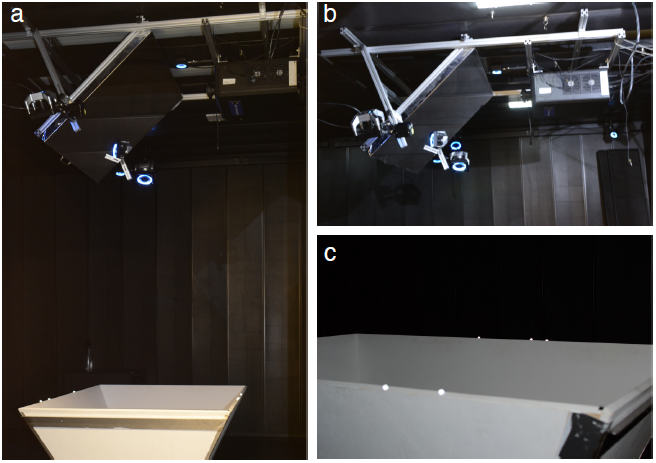
Photographs of the ratCAVE setup. (**a**) Photograph of the full system showing arena, projector, mirror, cameras (our system uses twelve cameras, arranged about the recording chamber and above the arena; only five (shown with blue lighted rings, normally turned off) are visible here). (**b**) Close-up on the projector, mirror, and cameras. (**c**) Close-up on the arena showing retro-reflective markers attached; the increased brightness of the markers is created in the photo by the camera's flash, and is brightened during VR sessions by the infra-red lighting of the tracking cameras. This increased brightness is beyond the visual spectrum of rodent vision.

**Supplementary Figure 5.**
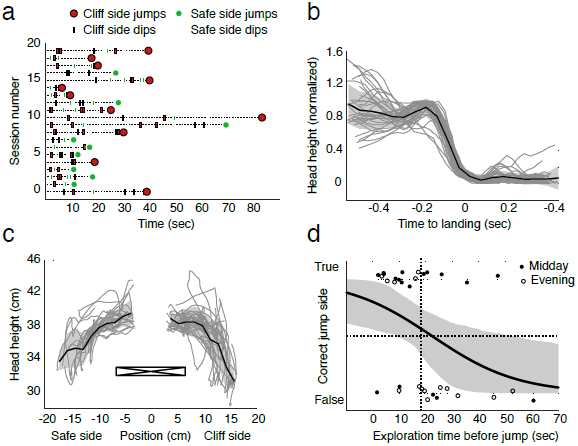
Visual cliff behavior trajectories. (**a**) Head-dipping and jumping events for each session. Rodents peeked over the edge of the board several times before choosing a side to jump off (example shown in Figure 2b), demonstrating risk assessment or exploration behavior. Head-dip events (rectangle markers) and jumps (circles) over time for each session. (**b**) Jump trajectories. Temporal dynamics of the rat’s head height (scaled by maximum height of jump over time relative to landing) is plotted as a function of time centered on the landing time. Individual jumps (gray lines), mean (black line) and standard deviation (gray shadow) are shown. (**c**) Head-dipping trajectories during rats’ visual inspection of the arena floors from the board (board cross-section shown in black, x-axis flipped with cliff on right side). We found no significant relationship between statistics of the head-dips and decision side. (**d**) Factors affecting jump side preference. Jump decision as a function of exploration time before jump. Note the difference in accuracy between first and second sessions recorded each day. Logistic regression found a significant correlation between exploration time and safe side preference (b = -.06, p < .05, solid black line, 68% CI as gray shading). Evening sessions with jump latencies greater than 18 secs (7 out of 33 sessions) were, as a result, excluded from the analysis of jump preference.

**Supplementary Figure 6.**
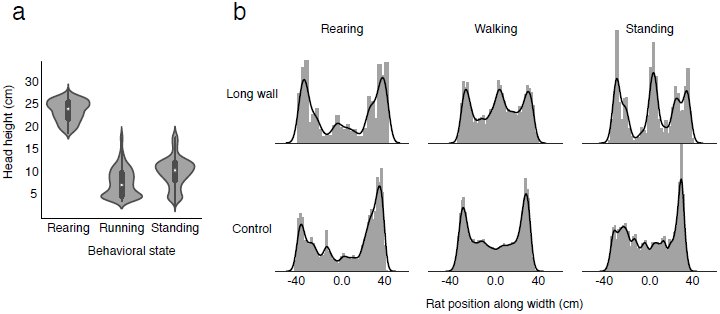
Virtual wall interaction. **(a)** Distribution of head height across different behavioral states (see Online Methods). **(b)** Occupancy along the arena width across different behaviors, in control and virtual wall conditions. Note that increased occupancy around the virtual wall location (arena center) occurs only when the virtual wall was present.

**Supplementary Figure 7.**
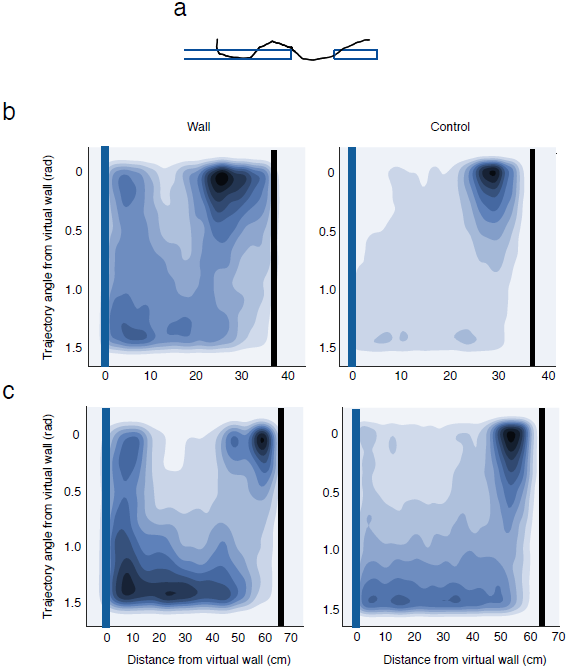
Thigmotaxis-like behavior along the virtual walls. (**a**) Example of a long trajectory parallel to the virtual wall. (**b**) Joint probability density of the orientation of locomotion trajectories with respect to the virtual wall (y axis, 0 rad means parallel to virtual wall) and distance to the location of the long virtual wall along arena width (x-axis) for sessions with the wall (left) and control sessions without a wall (right). Location of virtual (blue) and real arena (black) walls is indicated by thick lines on the plot. (**c**) Same as b for the short wall. Note the mode at 0 radians close to the wall (<10cm from virtual wall), resembling thigmotaxis behavior near the real walls for both conditions. The mode close to 1.5 rad corresponds both to trajectories that “deflect from” and those that “cross” the virtual wall (Fig. 3).

**Supplementary Figure 8.**
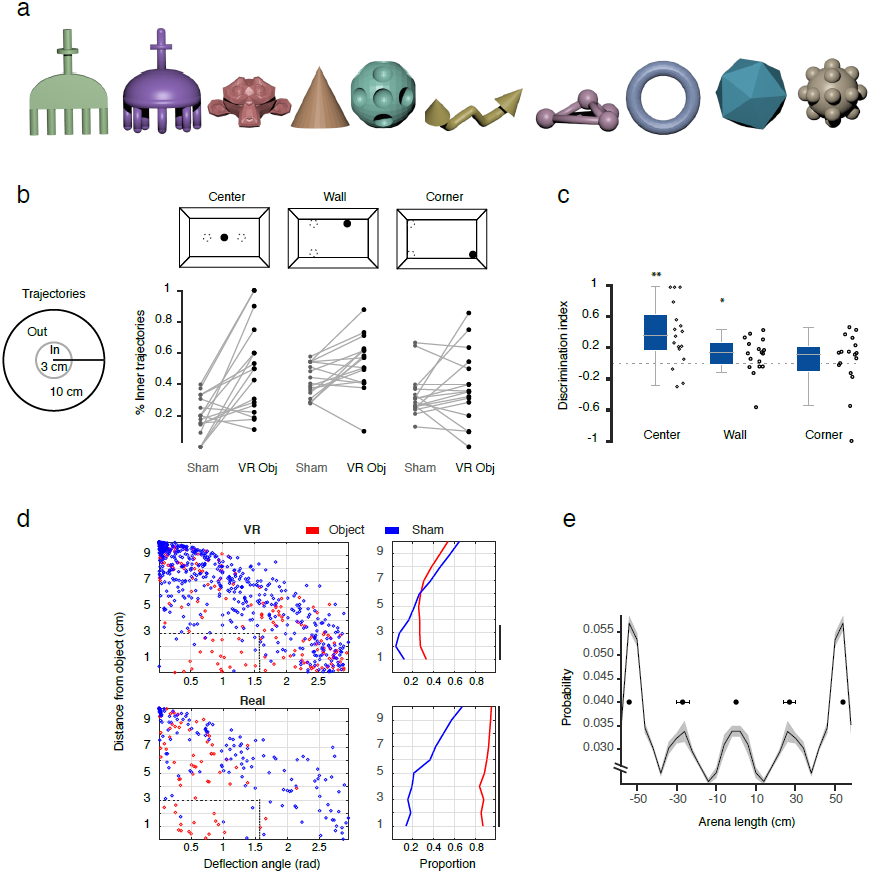
Object exploration analysis. (**a**) Virtual objects used in the task. Object shape and coloring were randomized between sessions. (**b**) Fine direction of trajectories towards virtual objects. Diagram on the left shows the areas used to calculate fine trajectories: percentage between inner (3 cm) and outer (10 cm) radius from the center of objects where consider as trajectories. Sham (dotted circles) and virtual object locations (filled circles) used in the calculations are shown for all objects (top). Percentage of inner trajectories for sham (Sham) and virtual object (VR Obj) locations (**c**) Discriminatory index from same data shown in b, significant discrimination index is observed only in center and wall object (Center: Z=2.889, n=17, ^**^ p<.01; Wall: Z=2.130, n=17, ^*^ p<.05; Corner Z=1.008, n=17, p=.08, Wilcoxon sign-rank test). (**d**) Deflection analysis for trajectories around the object. Scatter plots showing the relationship of arc angle made by trajectories entering and leaving the 10 cm circle around the object (deflection angle) and shortest distance between the trajectory to the object, for virtual (“VR”, top) and real (“Real”, bottom) objects for sham locations (blue) and object locations (red). Note that there is a concentration of trajectories in the vicinity of the object (~3 cm) with low arc angle value (<1.56 rad, 90 degrees), indicated by a dotted rectangle. These trajectories were used in the analysis in Fig. 4D, as they corresponded to trajectories that deflected from the object. Right column, cumulative proportion of deflecting trajectories (<1.56 rad) as function of distance from the object. Distances showing significant difference between object and sham conditions are indicated by thick black line on the right (p<.05, p-value calculated from a permutation distribution shuffled between conditions). (**e**) Example of the occupancy (projected on arena length) distribution of empty arena for the trials that followed object exploration trials, note the peaks that appear at the center of the arena (0, location of the center object), +/-30cm (location of the wall object), and +/-50 cm (location of the corner object), indicating a bias of rat behavior by potential spatial memory of the previous virtual object experience in these locations. Error bars above the modes at 0, 30 and 50cm show bootstrap confidence intervals of the mode position computed across control sessions.

